# Genome-wide Prediction of Small Molecule Binding to Remote Orphan Proteins Using Distilled Sequence Alignment Embedding

**DOI:** 10.1101/2020.08.04.236729

**Authors:** Tian Cai, Hansaim Lim, Kyra Alyssa Abbu, Yue Qiu, Ruth Nussinov, Lei Xie

## Abstract

Endogenous or surrogate ligands of a vast number of proteins remain unknown. Identification of small molecules that bind to these orphan proteins will not only shed new light into their biological functions but also provide new opportunities for drug discovery. Deep learning plays an increasing role in the prediction of chemical-protein interactions, but it faces several challenges in protein deorphanization. Bioassay data are highly biased to certain proteins, making it difficult to train a generalizable machine learning model for the proteins that are dissimilar from the ones in the training data set. Pre-training offers a general solution to improving the model generalization, but needs incorporation of domain knowledge and customization of task-specific supervised learning. To address these challenges, we develop a novel protein pre-training method, DIstilled Sequence Alignment Embedding (DISAE), and a module-based fine-tuning strategy for the protein deorphanization. In the benchmark studies, DISAE significantly improves the generalizability and outperforms the state-of-the-art methods with a large margin. The interpretability analysis of pre-trained model suggests that it learns biologically meaningful information. We further use DISAE to assign ligands to 649 human orphan G-Protein Coupled Receptors (GPCRs) and to cluster the human GPCRome by integrating their phylogenetic and ligand relationships. The promising results of DISAE open an avenue for exploring the chemical landscape of entire sequenced genomes.

## 1 Introduction

In spite of tremendous advances in genomics, the function of a vast number of proteins, particularly, their ligands are largely unknown. Among approximately 3,000 druggable genes that encode human proteins, only 5% to 10% of them have been targeted by an FDA-approved drug [1]. The proteins that miss ligand information are labeled as orphan proteins. The deorphanization of such proteins in the human and non-human genomes will not only shed a new light into their physiological or pathological roles but also provide new opportunities in drug discovery and precision medicine [2]. Many experimental approaches have been developed to deorphanize the orphan proteins such as G-protein coupled receptors (GPCRs) [3]. However, they are costly and time-consuming. Computational approaches may provide an efficient solution to generate testable hypotheses. With the increasing availability of solved crystal structures of proteins, homology modeling and protein-ligand docking are major methods for the effort in deorphanization [4]. However, the quality of homology models significantly deteriorates when the sequence identity between target and template is low. When a high-quality homology model is unavailable, structure-based methods, either physics-based or machine learning-based [5] could be unfruitful. Moreover, protein-ligand docking suffers from a high rate of false positives because it is incapable of accurately modeling conformational dynamics, solvation effect, crystalized water molecules, and other physical phenomena. Thus, it is tempting to develop new sequence-based machine learning approaches to the deorphanization, especially for the proteins that are not close homologs of or even evolutionarily unrelated to the ones with solved crystal structures or known ligands. Many machine learning methods have been developed to predict chemical-protein interactions (CPIs) from protein sequences. Early works [6] relied on feature engineering to create a representation of protein and ligand then use classifiers such as support vector machine to make final prediction. End-to-end deep learning approaches have recently gained a momentum [7][8]. In terms of the representation of protein sequence, Convolutional Neural Network (CNN) was applied in DeepDTA [9] and DeepConvDTI [10]. DeepAffinity [11] encoded a protein by its protein structure properties and used seq2seq [12] for the protein embedding. Most recently, TransformerCPI [13] used self-attention based Transformer to learn the CPI interaction, and achieved the state-of-the-art performance. Several other works incorporated additional domain knowledge for the protein representation. For example, WideDTI [14] utilized two extra features: protein motifs and domains (PDM). Quasi-Visual [5] proposed to use distance map from protein structures.

The aforementioned works mainly focused on filling in missing CPIs for the existing drug targets. One of the biggest challenges in these methods is that their performance could be suboptimal if the protein of interest is significantly different from those used in the training. Innovation is needed from the algorithm perspective for better protein representation learning. This motivated our proposed protein representation method, DIstilled Sequence Alignment Embedding (DISAE), for the purpose of deorphanization of remote orphan proteins. Although the number of proteins that have known ligands is limited, pre-trained language models such as ALBERT [15] open the door to incorporate the information of remote orphan proteins. Through the learning of functionally conserved information about protein families that span a wide range of protein space, DISAE is empowered to solve not only the problem of drug-target interaction prediction but also the challenge of deorphanization.

The major contributions of DISAE include distilled protein sequence self-supervised learning as well as optimized supervised fine-tuning. This novel pre-training-fine-tuning approach allows us to capture relationships between dissimilar proteins. Several studies have adopted the pre-training strategy to model protein sequences [16][17][18]. Different from existing methods that use original protein sequences as input, we distill the original sequence into an ordered list of triplets by extracting evolutionarily important positions from a multiple sequence alignment including insertions and deletions. Furthermore, we devise a module-based pre-training-fine-tuning strategy using ALBERT [15]. In the benchmark study, DISAE significantly outperforms other state-of-the-art methods for the prediction of ligand binding to dissimilar orphan proteins. We apply DISAE to the deorphanization of G-protein coupled receptors (GPCRs). GPCRs play a pivotal role in numerous physiological and pathological processes. Due to their associations with many human diseases and high druggabilities, GPCRs are the most studied drug targets [19]. Around one-third of FDA-approved drugs target GPCRs [19]. In spite of intensive studies in GPCRs, the endogenous and surrogate ligands of a large number of GPCRs remain unknown [3]. Using DISAE, we can confidently assign 649 orphan GPCRs in Pfam with at least one ligand. 106 of the orphan GPCRs find at least one approved GPCR-targeted drugs as ligand with estimated false positive rate lower than 0.05. These predictions merit further experimental validations. In addition, we cluster the human GPCRome by integrating their sequence and ligand relationships. The promising results of DISAE open the revenue for exploring the chemical landscape of all sequenced genomes. The code of DISAE is available from github^1^.

## 2 Methods

### 2.1 Protein sentence representation from distilled sequence alignment

A protein sequence is converted into an ordered list of amino acid fragments (words) with the following steps, as illustrated in Figure 1A.

**Figure 1:**
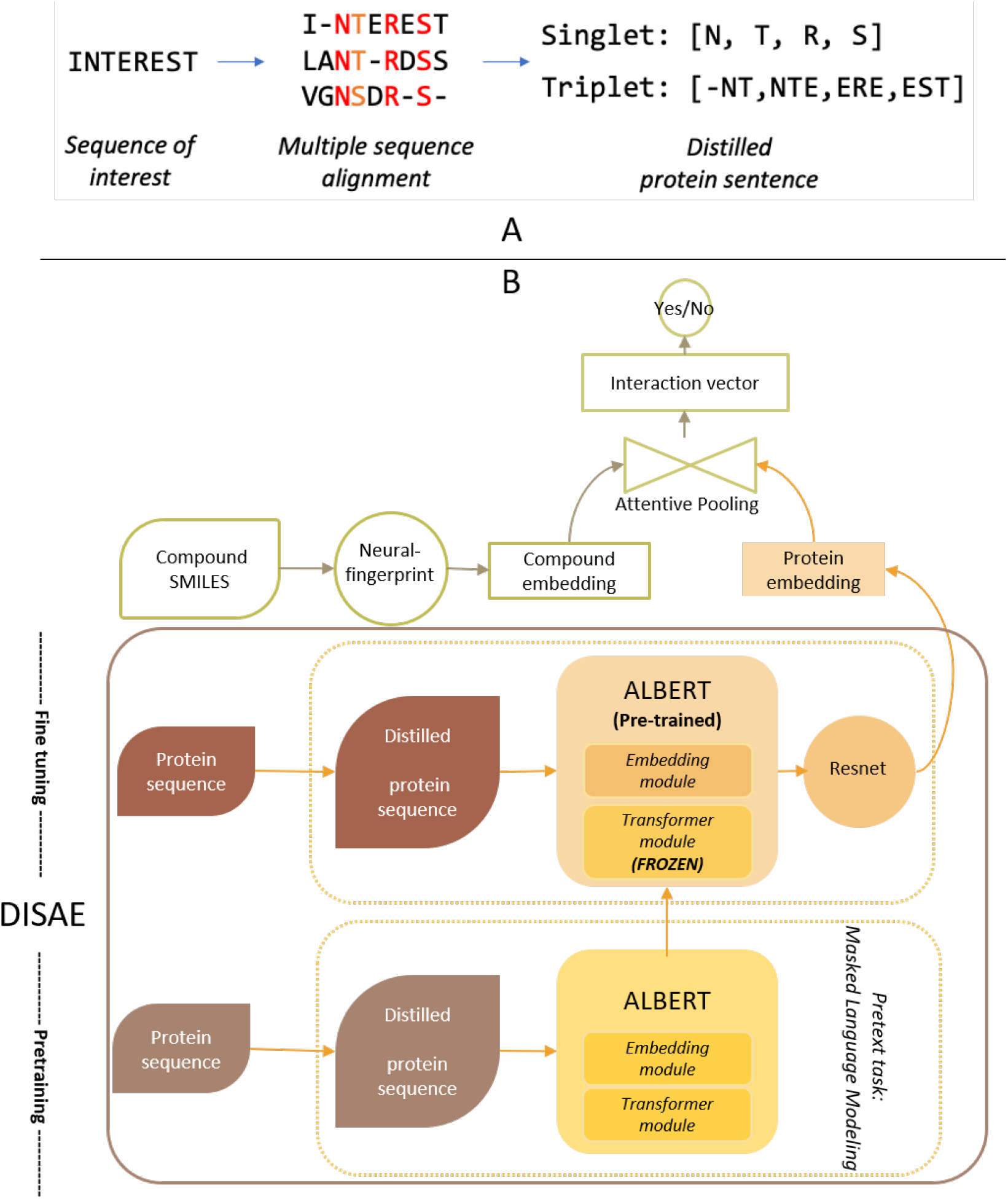
(A) Illustration of protein sequence representation from distilled sequence alignment. The high and medium conserved positions are marked as red and orange, respectively. (B) Architecture of deep learning model for the CPI prediction

1. Given a sequence of interest *S*, a multiple sequence alignment is constructed by a group of similar sequences to *S*. In this paper, the pre-computed alignments in Pfam [20] are used.
2. Amino acid conservation at each position is determined.
3. Starting from the most conserved positions, a pre-defined number of positions in the alignment are selected. In this study, the number of positions is set as 210. The length of positions was not optimized.
4. A word, either a single amino acid or a triplet of amino acids in the sequence, is selected at each position. The triplet may include gaps.
5. Finally, all selected words from the sequence form an ordered list following the sequence order, i.e. the sentence representation of the protein.

The rationale for the distilled multiple sequence alignment representation is to only use functionally or evolutionarily important residues and ignore others. It can be considered a feature selection step. In addition, the use of multiple sequence alignment will allow us to correlate the functional information with the position encoding. The distilled sequence will not only reduce the noise but also increase the efficiency in the model training since the memory and time complexity of language model is *O*(*n*^2^). It is noted that we use the conservation to select residues in a sequence because it is relevant to protein function and ligand binding. Other criteria (e.g., co-evolution) could be used depending on the down-stream applications (e.g., protein structure prediction).

### 2.2 Distilled sequence alignment embedding (DISAE) using ALBERT

It has been shown that natural language processing algorithms can be successfully used to extract biochemically meaningful vectors by pre-training BERT [21] on 86 billion amino acids (words) in 250 million protein sequences [22]. A recently released light version of BERT, called ALBERT [15], boasts significantly lighter memory uses with better or equivalent performances compared to BERT [21]. We extend the idea of unsupervised pre-training of proteins using ALBERT algorithm using distilled multiple sequence alignment representation. The distilled ordered list of triplets or singlets is used as the input for ALBERT pre-training. In this work, only masked language model (MLM) is used for the pre-training.

### 2.3 Architecture of deep learning model for CPI prediction

The deep learning model for the CPI prediction is mainly composed of three components, as shown in Figure 1B, protein embedding by DISAE, chemical compound embedding, and attention pooling with multilayer perceptron (MLP) to model CPIs. DISAE is described in the previous section. Note that once processed through ALBERT, each protein is represented as a matrix of size 210 by 312, where each triplet or singlet is a vector of length 312. During protein-ligand interaction prediction task, the protein embedding matrix is compressed using ResNet [23]. Once processed through ResNet layers, each protein is represented as a vector of length 256, which contains compressed information for the whole 210 input triplets or singlets for the corresponding protein.

Neural molecular fingerprint [24] is used for the chemical embedding. Small molecule is represented as a 2D graph, where vertices are atoms and edges are bonds. We use a popular graph convolutional neural network to process ligand molecules [25].

The attentive pooling is similar to the design in [8]. For each putative chemical-protein pair, the corresponding embedding vectors are fed to the attentive pooling layer, which in turn produces the interaction vector.

More details on the neural network model configurations can be found in Supplementary Table 1, 2, 3.

### 2.4 Module-based fine tuning strategy

When applying ALBERT to a supervised learning task, fine-tuning [15] is a critical step for the task-specific training following the pre-training. Pre-trained ALBERT already learned to generate protein representation in a meaningful way. However, it is also a design choice whether to allow ALBERT to get updated and trained together with the other components of the classification system during the fine-tuning [15][26]. Updating ALBERT during the fine-tuning will allow the pre-trained protein encoder to better capture knowledge from the training data while minimizing the risk of significant loss of knowledge obtained from the pre-training. Hence, to find the right balance, we experiment with ALBERT models that are partially unfrozen [26]. To be specific, major modules in the ALBERT model, embedding or transformer [27], are unfrozen separately as different variants of the model. The idea of “unfrozen” layers is widely used in Natural Language Processing (NLP), e.g., ULMFiT [26] where model consists of several layers. As training proceeds, layers are gradually unfrozen to learn the task-specific knowledge while safeguarding knowledge gained from pre-training. However, this is not straightforwardly applicable to ALBERT because ALBERT is not a linearly layered-up architecture. Hence, we apply a module-based unfrozen strategy.

### 2.5 Experiments Design

The purpose of this study is to build a model to predict chemical binding to novel orphan proteins. Therefore, we design experiments to examine the model generalization capability to the data not only unseen but also from significantly dissimilar proteins. We split training/validation/testing data sets to assess the performance of algorithms in three scenarios: 1. The proteins in the testing data set are significantly different from those in the training and validation data set based on the sequence similarity. 2. The ligands in the testing data are from a different gene family from that in the training/validation data. 3. Whole data set is randomly split similar to most of the existing work.

To examine the effect of different pre-training-fine-tuning algorithms, we organize experiments in three categories of comparison, as shown in Table 1:

a. THE EFFECT OF DISTILLED SEQUENCE: We assess whether distilled sequences improve testing performance by comparing ALBERT models where all configurations are the same except for sequence being distilled or not.
b. THE EFFECT OF VOCABULARY: Taking protein sequence as a sentence, its vocabulary could be built in many ways. We compare taking the singlet as vocabulary against the triplet as vocabulary.
c. THE EFFECT OF PRE-TRAINING: We assess how unlabeled protein sequences affect the performance of the classification. We compare ALBERT pre-trainined on whole Pfam alone against one pre-trainined on GPCRs alone and one without pre-training.
d. THE EFFECT OF FINE-TUNING: We compare three ALBERT models: ALBERT all unfrozen, ALBERT frozen embedding, and ALBERT frozen transformer [27]. All of these modes are pre-trained on the whole Pfam.

**Table 1:**
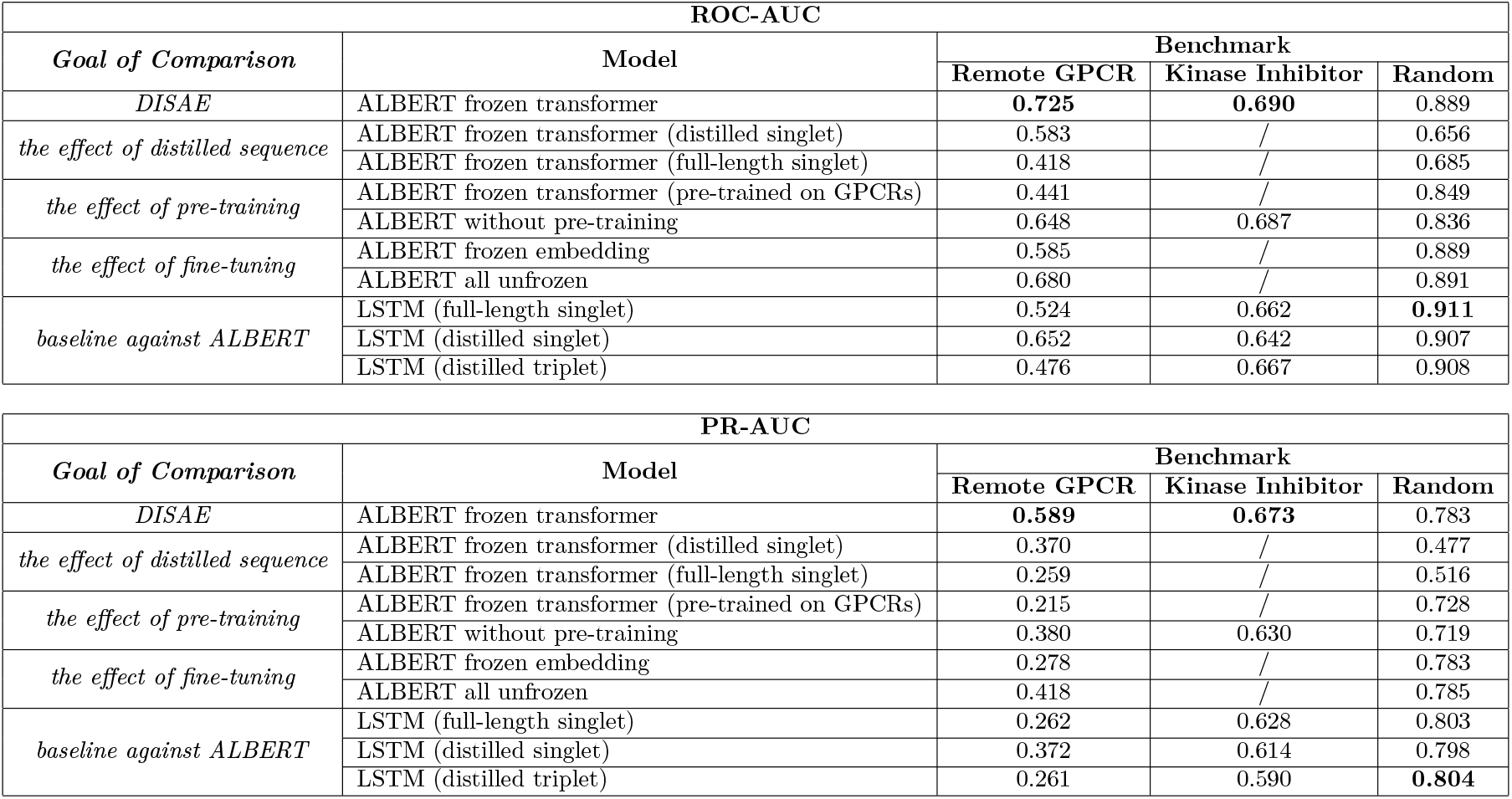
Test set performance under three data settings evaluated in ROC-AUC and PR-AUC. ALBERT pre-trained transformer-frozen model outperforms other models, and its performance is stable across all settings. Hence, it is recommended as the best model for the deorphanization. Six variants of DISAE models are compared to the frozen transformer one. Unless specified in the parentheses, ALBERT is pre-trained on whole Pfam proteins in the form of distilled triplets. The six DISAE variants are organized into three groups based on the goal of comparison. Protein similarity based splitting uses a threshold of bitscore 0.035

We use LSTM [28] and Transformer (equivalent to ALBERT without pre-training) as baselines. Three variants of LSTM models are tested to compare with the above three groups of experiments: LSTM with distilled triplets, LSTM with distilled singlets and LSTM with full-length singlets.

### 2.6 Dataset

Our task is to learn from large-scale CPI data to predict unexplored interactions. The quality and quantity of the training samples are critical for biologically meaningful predictions. Despite continuous efforts in the community, a single data source typically curates an incomplete list of our knowledge in protein-ligand activities. Thus, we integrated multiple high-quality, large-scale, publicly available databases of known protein-ligand activities. We extracted, split, and represented protein-ligand activity samples for training and evaluation of machine learning-based predictions.

#### 2.6.1 Sequence data for ALBERT pre-training

Proteins sequences are first collected from Pfam-A [20] database. Then, sequences are clustered by 90% sequence identity, and a representative sequence is selected from each cluster. The filtered non-redundant 35181 sequences are used for the ALBERT pre-training. To construct protein sentences from the distilled sequence alignment, the original alignment and conservation score from each Pfam-A family are used. As a comparison, the sequences only from GPCR family are used to pre-train a separated ALBERT model.

We used Pfam alignments directly. From the consensus alignment sequence, we collected positions of high-confidence and low-confidence conserved amino acids together with conservatively substituted ones. We picked these positions from each of the target GPCRs. As a result, each GPCR is represented by 210 amino acids, which may contain gaps. The 210 amino acids are then treated as a sentence of triplets or singlets. The triplets or singlets are used to train the ALBERT model with the following parameters: maximum sequence length = 256, maximum predictions per sentence = 40, word masking probability = 0.15, and duplication factor = 10. Note that the order of different GPCRs in multiple sequence alignment may not be biologically meaningful. Thus, we did not apply the next sentence prediction task during the pre-training.

#### 2.6.2 Binding assay data for supervised learning

We integrated protein-ligand activities involving any GPCRs from ChEMBL [29] (ver. 25), BindingDB [30] (downloaded Jan 9, 2019), GLASS [31] (downloaded Nov 26, 2019), and DrugBank [32] (ver. 5.1.4). Note that BindingDB also contains samples drawn from multiple sources, including PubChem, PDSP *K*_*i*_, and U.S. Patent. From ChEMBL, BindingDB, and GLASS databases, protein-ligand activity assays measured in three different unit types, *pK_d_*, *pK_i_*, and *pIC*_50_ are collected. Log-transformation were performed for activities reported in *K*_*d*_, *K*_*i*_, or *IC*_50_. For consistency, we did not convert different activity types. For instance, activities reported in *IC*50 are converted only to *pIC*_50_, but not any other activity types. The activities in log-scale were then binarized based on the thresholds of activity values. Protein-ligand pairs were considered active if *pIC*_50_ < 5.3, or *pK*_*d*_ or *pK*_*i*_ < 7.3, and inactive if *pIC*_50_ < 5.0, *pK*_*d*_ < 7.0 or *pK*_*i*_ < 7.0, respectively.

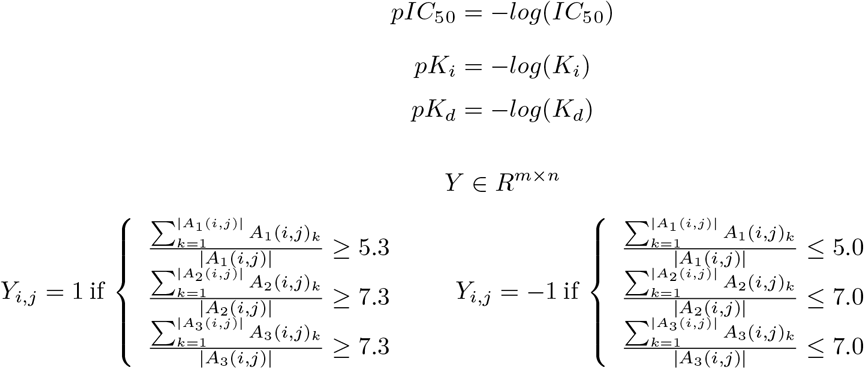

In the above equations, *m* and *n* are the total number of unique proteins and ligands, respectively. *A*_1_(*i, j*), *A*_2_(*i, j*), and *A*_3_(*i, j*) are the list of all activity values for the *i*th protein and *j*th ligand in *pIC*_50_, *pK*_*d*_, and *pK*_*i*_, respectively. |*A*| denotes the cardinality of the set *A*. Note that there are gray areas in the activity thresholds. Protein-ligand pairs falling in the gray areas were considered undetermined and unused for training. If multiple activities were reported for a protein-ligand pair, their log-scaled activity values were averaged for each activity type and binarized accordingly. In addition, we collected active protein-ligand associations from DrugBank and integrated with the binarized activities mentioned above. Inconsistent activities (e.g. protein-ligand pairs that appear both active and inactive) were removed. There are total 9705 active and 25175 inactive pairs, respectively, in the benchmark set.

To test the model generalization capability, a protein similarity based data splitting strategy is implemented. First, pair-wise protein similarity expressed by bit-score is calculated using BLAST [33] for all GPCRs in the benchmark. Then, according to the similarity distribution, a similarity threshold is set for the splitting. The bit-score threshold is 0.035 in this study. The sequences are clustered such that the sequences in the testing set are significantly dissimilar to those in the training/validation set. After the splitting, there are 25114, 6278, and 3488 samples for training, validation, and testing, respectively. Distribution of protein sequence similarity scores can be found in Supplementary Figure 1.

### 2.7 Ensemble model for the GPCR deorphanization

All annotated GPCR-ligand binding pairs are used to build prediction model for the GPCR deorphanization. To reduce possible overfitting, an ensemble model is constructed using the chosen ALBERT model. Following the strategy of cross-validation [34], three DISAE models are trained. Similar to the benchmark experiments, a hold-out set is selected based on protein similarity, and is used for the early stopping at the preferred epoch for each individual model. Max-voting [35] is invoked to make final prediction for the orphan human GPCRs.

To estimate the false positive rate of predictions on the orphan-chemical pairs, a prediction score distribution is collected for known positive pairs and negative pairs of testing data. Assuming that the prediction score of orphan pairs have the same distribution as that of the testing data, for each prediction score, a false positive rate can be estimated based on the score distribution of true positives and negatives.

### 2.8 SHAP analysis

Kernel SHAP [36] is used to calculate SHAP values for distilled protein sequences. It is a specially-weighted local linear regression to estimate SHAP values for any model. The model used is the classifier that fine-tunes the ALBERT frozen transformer, and is pre-trained on whole Pfam in distilled triplets under the remote GPCR setting. The data used is the testing set generated under the same remote GRPC setting. Although the whole input features to the classifier consist of both distilled protein sequence and chemical neural fingerprint, only protein feature is of the interest. Hence, when calculating the base value (a value that would be predicted if we do not know any features) required by SHAP analysis, all testing set protein sequence are masked with the same token, while chemical neural fingerprint remain untouched. Therefore, the base value is the average prediction score without protein feature and solely relying on chemical feature. Since the distilled sequences are all set to be of length 210, the SHAP values are the feature importance for each of the 210 positions.

### 2.9 Hierarchical clustering of the human GPCRome

The pairwise distance between two proteins is determined by the cosine similarity of the DISAE embedding vector after pre-training-fine-tuning, which incoporates both sequence and ligand information. R pacakge “ape” (Analyses of Phylogenetics and Evolution) [37] and treeio [38] is used to convert a distance matrix of proteins into a Newick tree format. Gtree [39][40] package is used to plot the tree in a circular layout.

## 3 Results and Discussion

### 3.1 Pre-trained DISAE triplet vector is biochemically meaningful

When pre-training ALBERT model with masked triplets of proteins, the masked word prediction accuracy reached 0.982 and 0.984 at the 80,000th and the 100,000th training step, respectively. For comparison, the pre-training accuracy when using BERT [21] was 0.955 at 200,000th training step. It may be because a larger batch size can be used in ALBERT than in BERT. The distilled sequence alignment embedding (DISAE) from ALBERT pre-training is subsequently used for the CPI prediction. To evaluate whether the pre-trained DISAE vector is biochemically meaningful, we extracted the pre-trained DISAE vector for all possible triplets. We then used t-SNE to project the triplet vector in a 2D space. As shown in Supplementary Figure 2, the triplet vectors formed distinct clusters by the properties of amino acid side chains in the third amino acid of triplets, especially at level 3 and 4. Triplets containing any ambiguous or uncommon amino acids, such as amino acid U for selenocysteine or X for any unresolved amino acids, formed a large cluster that did not form a smaller group (large group of black dots in each scatter plot), suggesting that the information regarding such rare amino acids are scarce in the pre-training dataset. When there is no ambiguity in triplets, they form clearly separated clusters, meaning that the pre-training process indeed allows the model to extract biochemically meaningful feature vectors from the proteins as sentences. The same clustering trends were also observed when triplets are grouped by individual amino acids, rather than their physicochemical properties of side chains (Supplementary Figure 3).

### 3.2 DISAE significantly outperforms state-of-the-art models for predicting ligands of remote orphan proteins

When the proteins in the testing data are significantly dissimilar to those in the training data, the best model is ALBERT pre-trained transformer-frozen model, with a ROC-AUC of 0.725 and a PR-AUC of 0.589, respectively, as shown in Table 1 and Figure 2A and 2B. As a comparison, the ROC-AUC of ALBERT without the pre-training model and LSTM model is 0.648 and 0.476, respectively; and the PR-AUC of ALBERT without the pre-training model and LSTM model is 0.380 and 0.261, respectively. Furthermore, as shown in the insert of Figure 2A, ALBERT pre-trained transformer-frozen model not only has the highest ROC-AUC and PR-AUC but also far exceeds other models at the range of low false positive rates. For example, at the false positive rate of 0.05, the number of true positives detected by ALBERT pre-trained transformer-frozen model is almost 6 times more than that correctly predicted by ALBERT without the pre-training model or LSTM models when using the same sequence representation.

**Figure 2:**
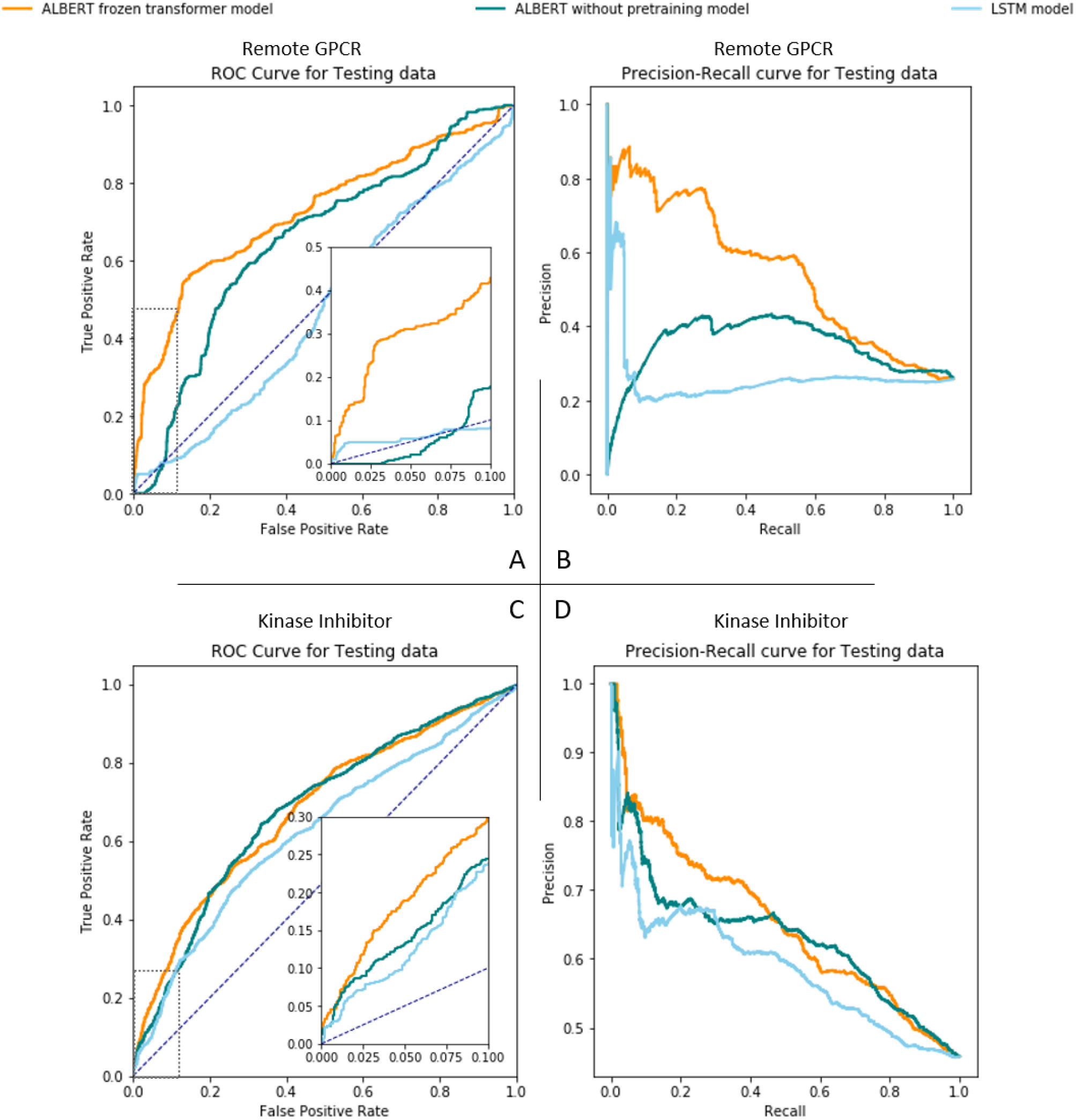
Performance comparison of ALBERT pre-trained transformer-frozen model, ALBERT without pre-training model, and LSTM model. **(A, B)**: ROC- and PR-curves for the prediction of ligand binding to remote GPCRs (bitscore<0.035). **(C, D)**: ROC- and PR-curves for the classification on testing set in the cross-gene-family kinase inhibitor benchmark. All the models are trained on distilled triplets.

When the train/test set is split at random, i.e. there are similar proteins in the testing set to those in the training set, the performance of LSTM models is slightly better than the ALBERT pre-trained models. However, the performance of the LSTM model significantly deteriorates when the proteins in the testing set are different from those in the training set. The ROC-AUC drops from over 0.9 to ~0.4-0.6, and PR-AUC decreases from over 0.8 to ~0.2-0.3, while ALBERT pre-trained models could still maintain the ROC-AUC ~0.7 and the PR-AUC ~0.5. These results suggest that the supervised learning alone is prone to overfitting. The pre-training of DISAE, which uses a large number of unlabeled samples improve the generalization power. The training curves in Supplementary Figure 4, 5, 6 further support that ALBERT frozen transformer is generalizable, thus can reliably maintain its high performance when used for the deorphanization. When evaluated by the dissimilar protein benchmark, the accuracy of training keeps increasing with the increased epochs; and the performance of ALBERT pre-trained transformer-frozen model is slightly worse than the most of models. However, the PR-AUC of ALBERT pre-trained transformer-frozen model is relatively stable and significantly higher than other models on the testing data.

### 3.3 DISAE significantly outperforms state-of-the-art models for predicting ligand binding promiscuity across gene families

We further evaluate the performance of different variants of ALBERT pre-trained and LSTM models on the prediction of ligand binding promiscuity across gene families. Specifically, we predict the binding of kinase inhibitors to GPCRs on the model that is trained using only GPCR sequences and their ligands that are not known kinase inhibitors. Although the kinase and the GPCRs belong to two completely different gene families in terms of sequence, structure, and function, a number of kinase inhibitors can bind to GPCRs as an off-target. Interestingly, although DISAE is not extensively trained using a comprehensive chemical data set, the ALBERT pre-trained transformer-frozen model outperforms the LSTM and the ALBERT without the pre-training. As shown in Figure 2C, 2D and Table 1, both sensitivity and specificity of the ALBERT pre-trained transformer-frozen model outperforms other models. The ROC-AUC of ALBERT pre-trained transfomer-frozen model, ALBERT without the pre-training, LSTM model is 0.690, 0.687, and 0.667, respectively. Their PR-AUC is 0.673, 0.630, and 0.590, respectively. The improvement is the most obvious in the region of low false positive ratio (the insert of Figure 2C). At the false positive rate of 0.05, the number of true positives detected by the ALBERT pre-trained transformer-frozen model is about 50% as many as that by Transfomer-like and LSTM models. This observation implies that the sequence pre-training captures certain ligand binding site information across gene families.

### 3.4 The effect of distilled sequence representation, pre-training, and fine-tuning

For the three major pre-training-fine-tuning configurations of DISAE, <u>*distilled triplets sequence representation*, *pre-trained on whole Pfam*</u>, and <u>*fine-tuned with frozen transformer*</u> (“ALBERT frozen transformer” in Table 1), are recommended as the best performed model for CPI prediction and derophanization tasks. This conclusion is reached through a series of controlled experiments as shown in Table 1 under the dissimilar protein and cross-gene-family benchmarks.

a. Distilled sequence is preferred over non-distilled sequence With the same pre-training on whole Pfam and fine-tuning on the frozen transformer, the ALBERT model using the distilled sequence clearly outperforms that using the full-length sequence as evaluated by both ROC-AUC and PR-AUC. It is true for the LSTM models as well.
b. Triplet is preferred over singlet on the ALBERT model Under the same pre-training and fine-tuning settings, both ROC-AUC and PR-AUC of the ALBERT pre-trained model that uses the distilled triplets are significantly higher than the those when the distilled singlet is used. However, for LSTM model, the distilled singlet outperforms the distilled triplet. It seems that the triplet encodes more relational information on the remote proteins than close homologs.
c. Pre-training on larger dataset is preferred With the same fine-tuning strategy, frozen transformer, and use of distilled triplets, the model that is pre-trained on whole Pfam performs better than that pre-trained on the GPCR family alone or without pre-training in terms of both ROC-AUC and PR-AUC.
d. Partial frozen transformer is preferred With the same pre-training on whole Pfam and fine-tuning on the distilled triplets, the ALBERT pre-trained transformer-frozen model outperforms all other models.

### 3.5 DISAE learns biologically meaningful information

Interpretation of deep learning is critical for its real-world applications. To understand if the trained DISAE model is biologically meaningful, we perform the model explainability analysis using SHapley Additive exPlanation (SHAP) [36]. SHAP is a game theoretic approach to explain the output of any machine learning model. Shapley values could be interpreted as feature importance. We utilize this tool to get a closer look into the internal decision making of DISAE’s prediction by calculating Shapley values of each triplet of a protein sequence. The average Shapley values of CPIs for a protein is used to highlight important positions for this protein.

Figure 3 shows the distribution of a number of residues of 5-hydroxytryptamine receptor 2B on its structure, which are among 21 (10%) residues with the highest SHAP values. Among them, 6 residues (T140, V208, M218, F341, L347, Y370) are located in the binding pocket. L378 is centered in the functional conserved NPxxY motif that connects the transmembrane helix 7 and the cytoplasmic helix 8 and plays a critical role in the activation of GPCRs [41][42]. P160 and I161 is the part of intracellular loop 2, while I192, G194, I195, and E196 are located in the extracellular loop 2. The intracellular loop 2 interacts with the P-loop of G-protein [43]. It is proposed that the extracellular loop 2 may play a role in the selective switch of ligand binding and determine ligand binding selectivity and efficacy [44][45][46][47]. The functional impact of other residues are unclear. Nevertheless, more than one half of top 21 residues ranked by SHAP values can explain the trained model. The enrichment of ligand binding site residues is statistically significant (p-value=0.01). These results suggest that the prediction from DISAE can provide biologically meaningful interpretations.

**Figure 3:**
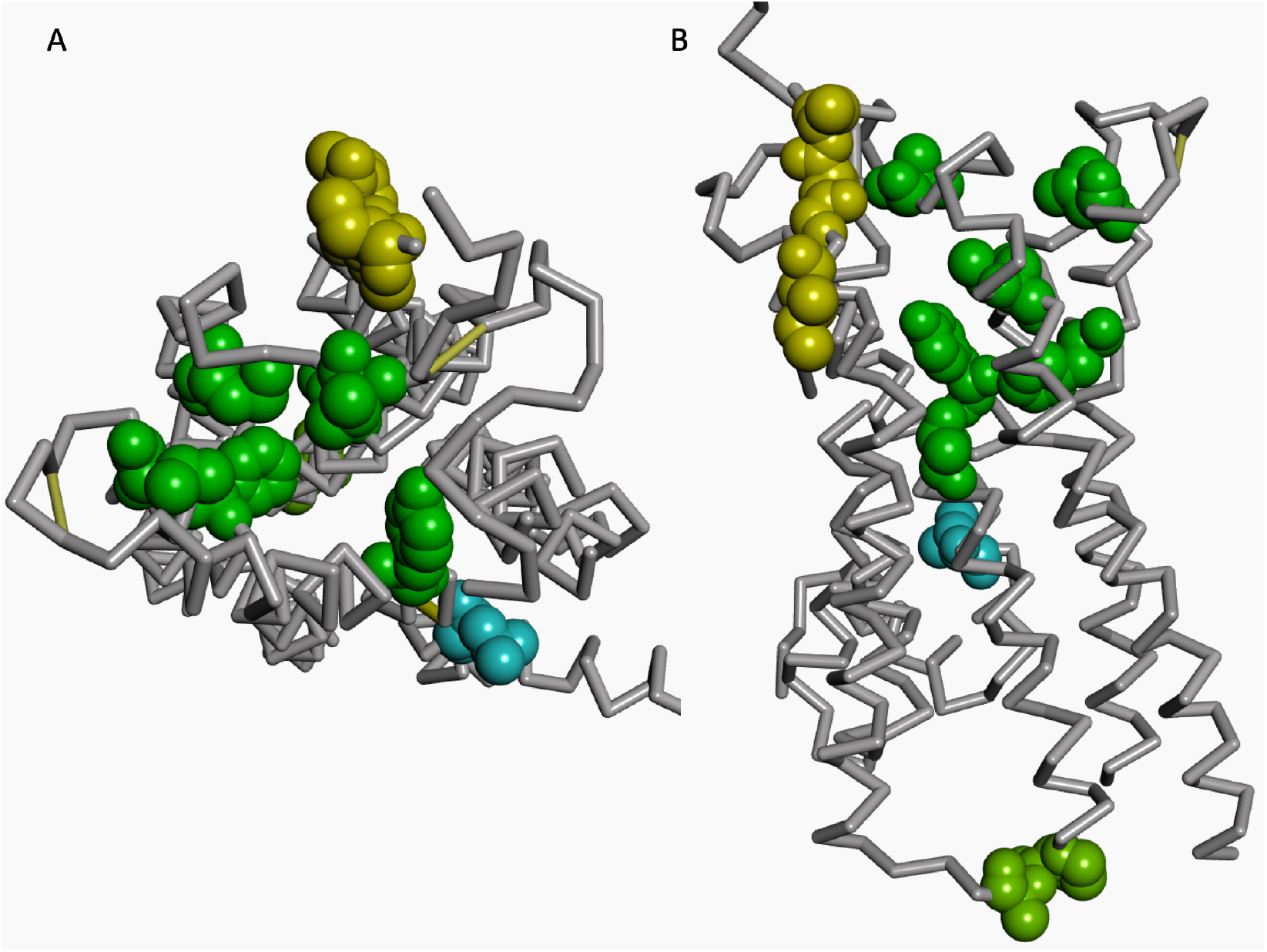
(A) top and (B) side view of structure of 5-hydroxytryptamine receptor 2B (UNIPROT id: 5HTB2_HUMAN, PDB ID: 4IB4). The residues among those with the top 21 ranked SHAP values are shown in dark green, blue, yellow, and light green colored CPK mode for the amino acids in the binding pocket, NPxxY motif, extracellular loop, and intracellular loop, respectively.

### 3.6 Application to the hierarchical classification and deorphanization of human GPCRs

With the established generalization power of DISAE, we use ***ALBERT transformer-frozen model** pre-trained on <u>whole Pfam</u> in <u>distilled triplets</u> form* to tackle the challenge of deorphanization of human GPCRs, due to its consistently excellent performance by all evaluation metrics in the benchmarks.

We define the orphan GPCRs as those which do not have known small molecule binders. 649 human GPCRs that are annotated in Pfam families PF00001, PF13853, and PF03402 are identified as the orphan receptors.

Studies have suggested that the classification of GPCRs should be inferred by combining sequence and ligand information [48]. The protein embedding of the DISAE model after pre-training-fine-tuning satisfies this requirement. Therefore, we use the cosine similarity between the embedded vector of protein as a metric to cluster the human GPCRome, which includes both non-orphan and orphan GPCRs. The hierarchical clustering of GPCRs in the Pfam PF00001 is shown in Figure 4. Supplementary Figure 7 and 8 show the hierarchical clustering of PF13853 and PF03402, respectively.

**Figure 4:**
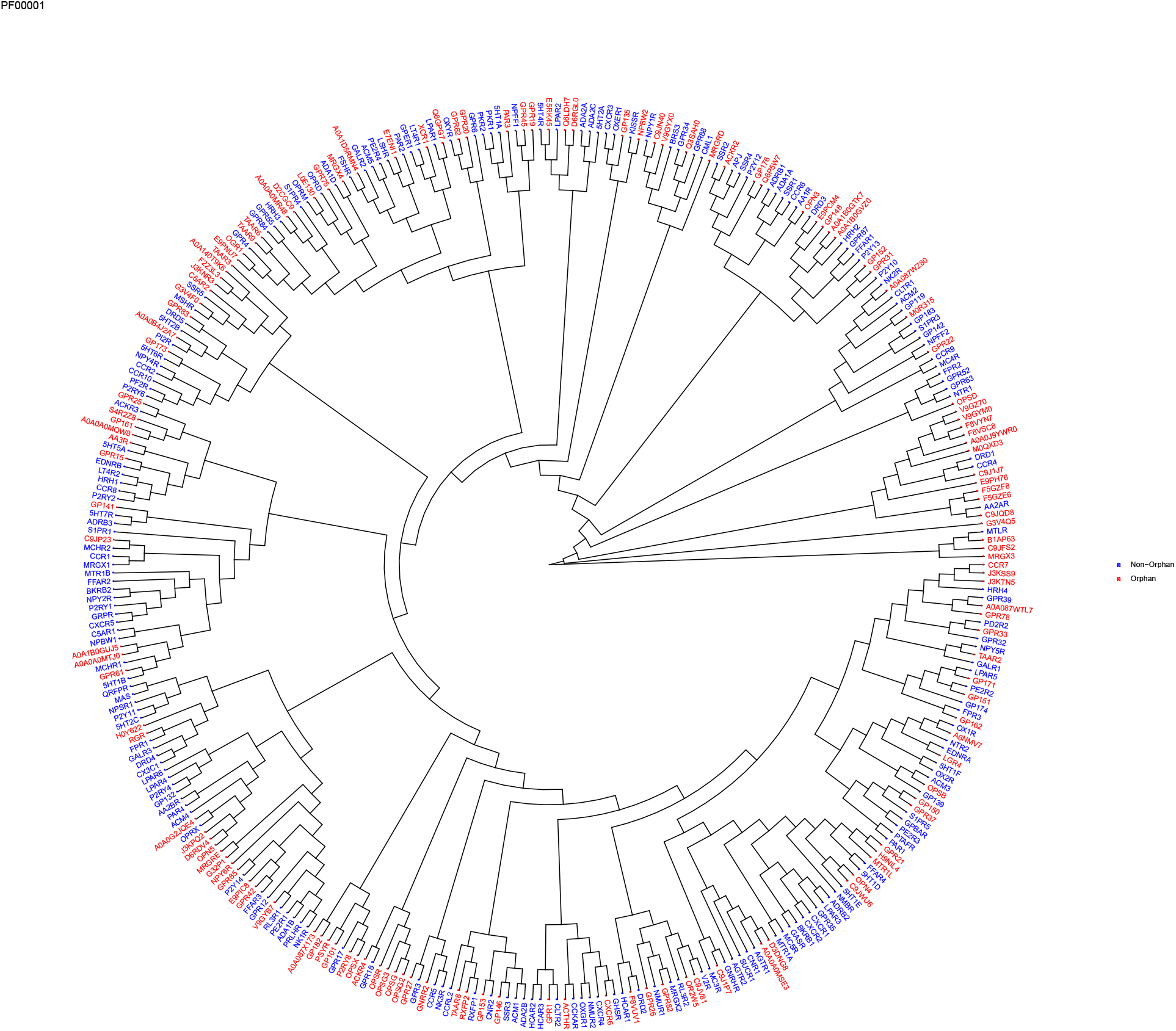
Hierarchical clustering of human GPCRs in the Pfam PF00001. Non-orphan and orphan GPCRs are labeled in blue and red, respectively

Table 2 provides examples of predicted approved-drug bindings to the orphan GPCRs by DISASE with high confidence (false positive rate < 5.0e-4). The complete list of orphans human GPCRs paired with 555 approved GPCR-targeted drugs is in the Supplementary Table 4. The predicted potential interactions between the approved drug and the orphan receptor will not only facilitate designing experiments to deorphanize GPCRs but also provide new insights into the mode of action of existing drugs for drug repurposing, polypharmacology, and side effect prediction.

**Table 2:** Example of deorphanization prediction from DISAE ensemble models.

| Orphan Receptor (Uniprot Id) | Pfam | Drug | Drug Target |
| --- | --- | --- | --- |
| A0A0C4DFX5 | PF13853 | Isoetharine | ADRB1, ADRB2 |
| A0A126GVR8 | PF13853 | Ganirelix | GNRHR |
| A0A1B0GTK7 | PF00001 | Levallorphan | OPRM1 |
| A0A1B0GVZ0 | PF00001 | Xamoterol | ADRB1, ADRB2 |
| A0A286YFH6 | PF13853 | Degarelix | GNRHR |
| A3KFT3 | PF13853 | Degarelix | GNRHR |
| C9J1J7 | PF00001 | Levallorphan | OPRM1 |
| C9JQD8 | PF00001 | Lixisenatide | GLP1R |
| E9PH76 | PF00001 | Ganirelix | GNRHR |
| E9PPJ8 | PF13853 | Xamoterol | ADRB1, ADRB2 |

## 4 Conclusion

Our primary goal in this paper is to address the challenge of predicting ligands of orphan proteins that are significantly dissimilar from proteins that have known ligands or solved structures. To address this challenge, we introduce new techniques for the protein sequence representation by the pre-training of distilled sequence alignments, and the module-based fine-tuning using labeled data. Our approach, DISAE, is inspired by the state-of-the-art algorithms in NLP. However, our results suggest that the direct adaption of NLP may be less fruitful. The successful application of NLP to biological problems requires the incorporation of domain knowledge in both pre-training and fine-tuning stages. In this regard, DISAE significantly improves the state-of-the-art in the deorphanization of dissimilar orphan proteins. Nevertheless, DISAE can be further improved in several aspects. First, more biological knowledge can be incorporated into the pre-training and fine-tuning at both molecular level (e.g. protein structure and ligand binding site information) and system level (e.g. protein-protein interaction network). Second, in the framework of self-supervised learning, a wide array of techniques can be adapted to address the problem of bias, sparsity, and noisiness in the training data. Put together, new machine learning algorithms that can predict endogenous or surrogate ligands of orphan proteins open up a new avenue for deciphering biological systems, drug discovery, and precision medicine.

## Acknowledgement

This project has been funded in whole or in part with federal funds from the National Institute of General Medical Sciences of National Institute of Health (R01GM122845), the National Institute on Aging of the National Institute of Health (R01AD057555), and the National Cancer Institute of National Institutes of Health, under contract HHSN261200800001E. The content of this publication does not necessarily reflect the views or policies of the Department of Health and Human Services, nor does mention of trade names, commercial products or organizations imply endorsement by the US Government. This Research was supported [in part] by the Intramural Research Program of the NIH, National Cancer Institute, Center for Cancer Research and the Intramural Research Program of the NIH Clinical Center.

## Footnotes

1 https://github.com/XieResearchGroup/DISAE

