## Supplementary Information for "Genome-wide Prediction of Small Molecule Binding to Remote Orphan Proteins Using Distilled Sequence Alignment Embedding"

July 2020

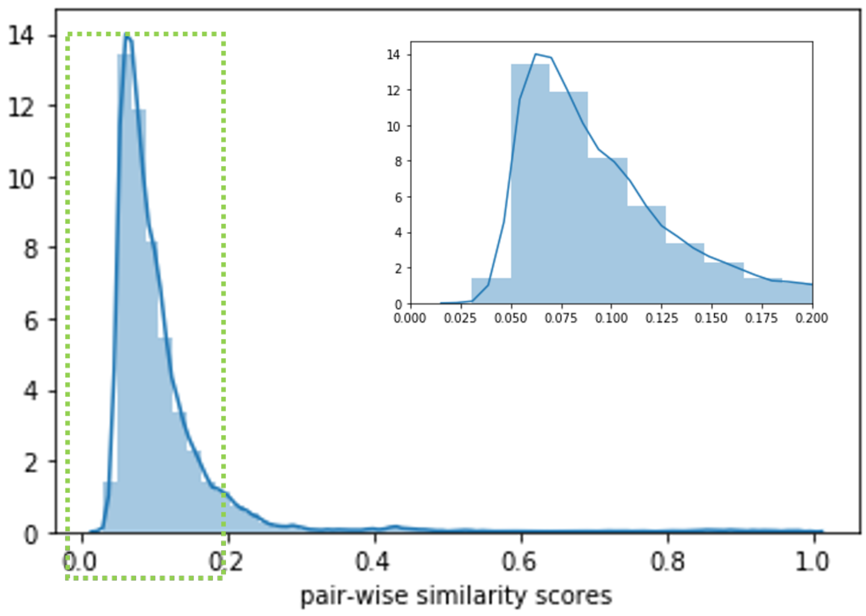


Supplementary Figure 1: Protein pair wise similarity distribution of benchmark data:To simulate realistic deorphanization scenario, where remote orphan protein could be significantly different from data used to train models, we design a protein similarity based data splitting strategy. First, pair wise bit-scores are calculated with standard BLAST+ package. A distribution could be found in Figure 1. Then, setting a threshold of 0.035, proteins with pair-wise similarity lower than the threshold are keep separately in two groups. The protein-chemical activity pairs with protein in the relative smaller group will be used as testing data. The left samples are then split into training and validation.

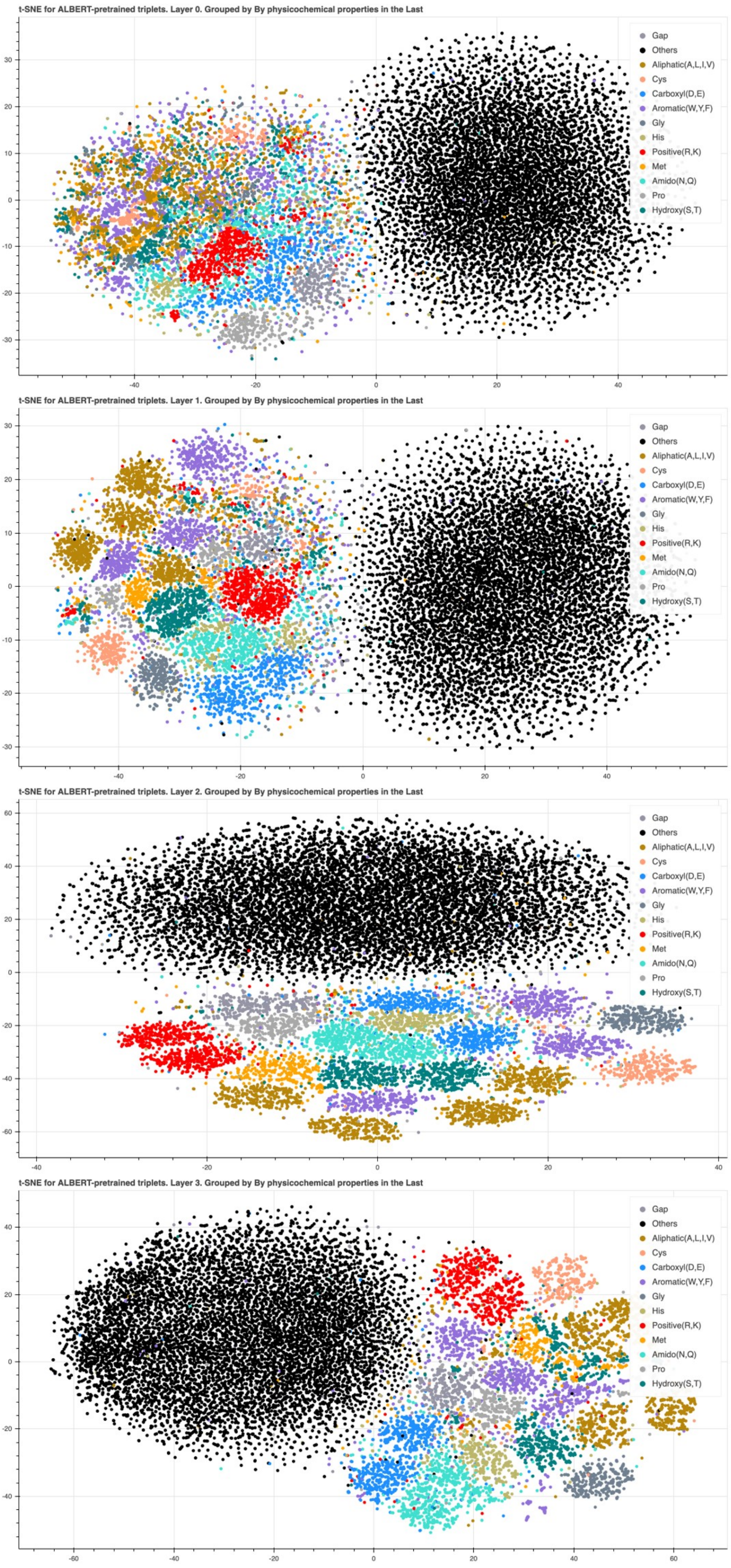


Supplementary Figure 2: Clustering of pre-trained triplet DISAE vectors at (A) Level 1, (B) Level 2, (C) Level 3, and (D) Level 4 of ALBERT. The triplet is colored by the physiochemical properties of the third amino acid.

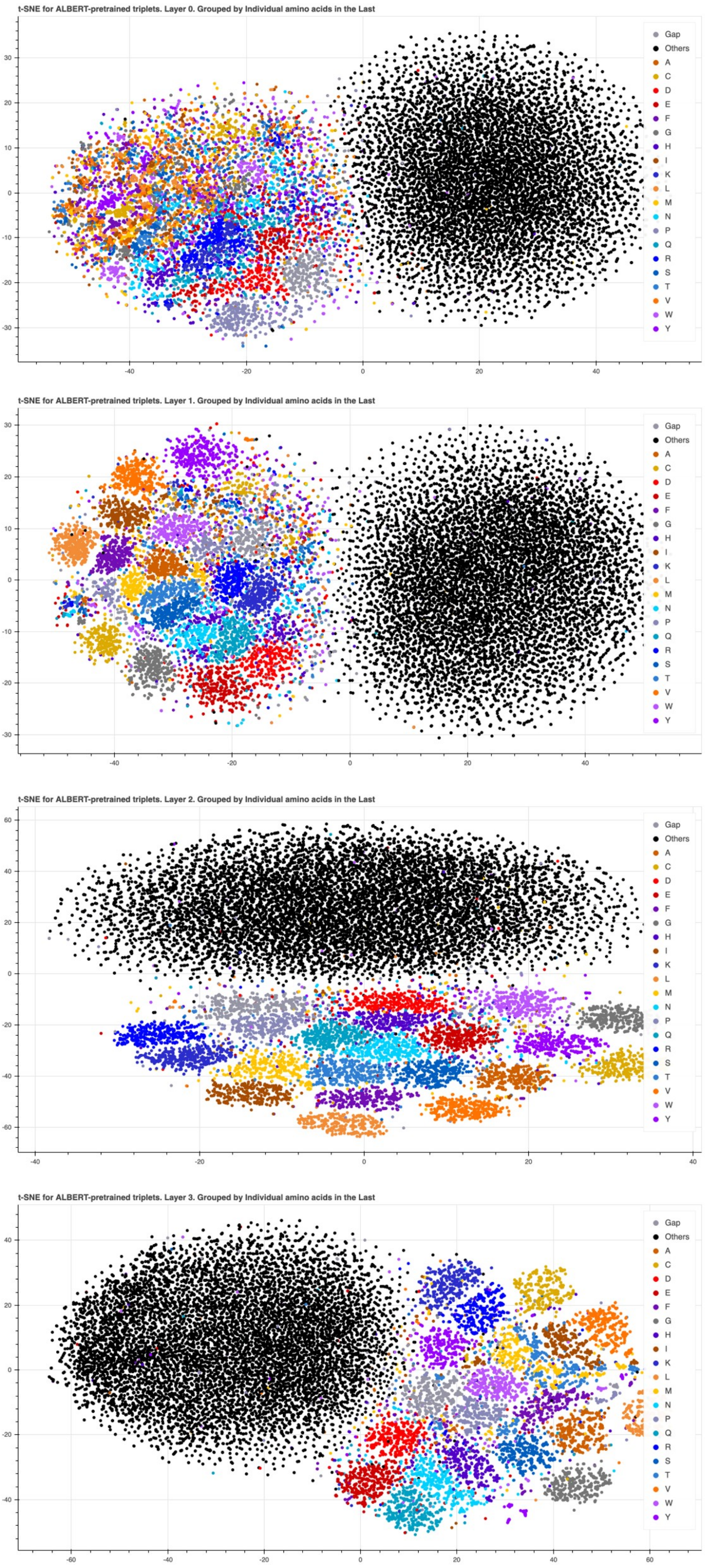


Supplementary Figure 3: The same clustering trends as in Supplementary Figure 2 were also observed when triplets are grouped by individual amino acid types, rather than their physicochemical properties of side chains.

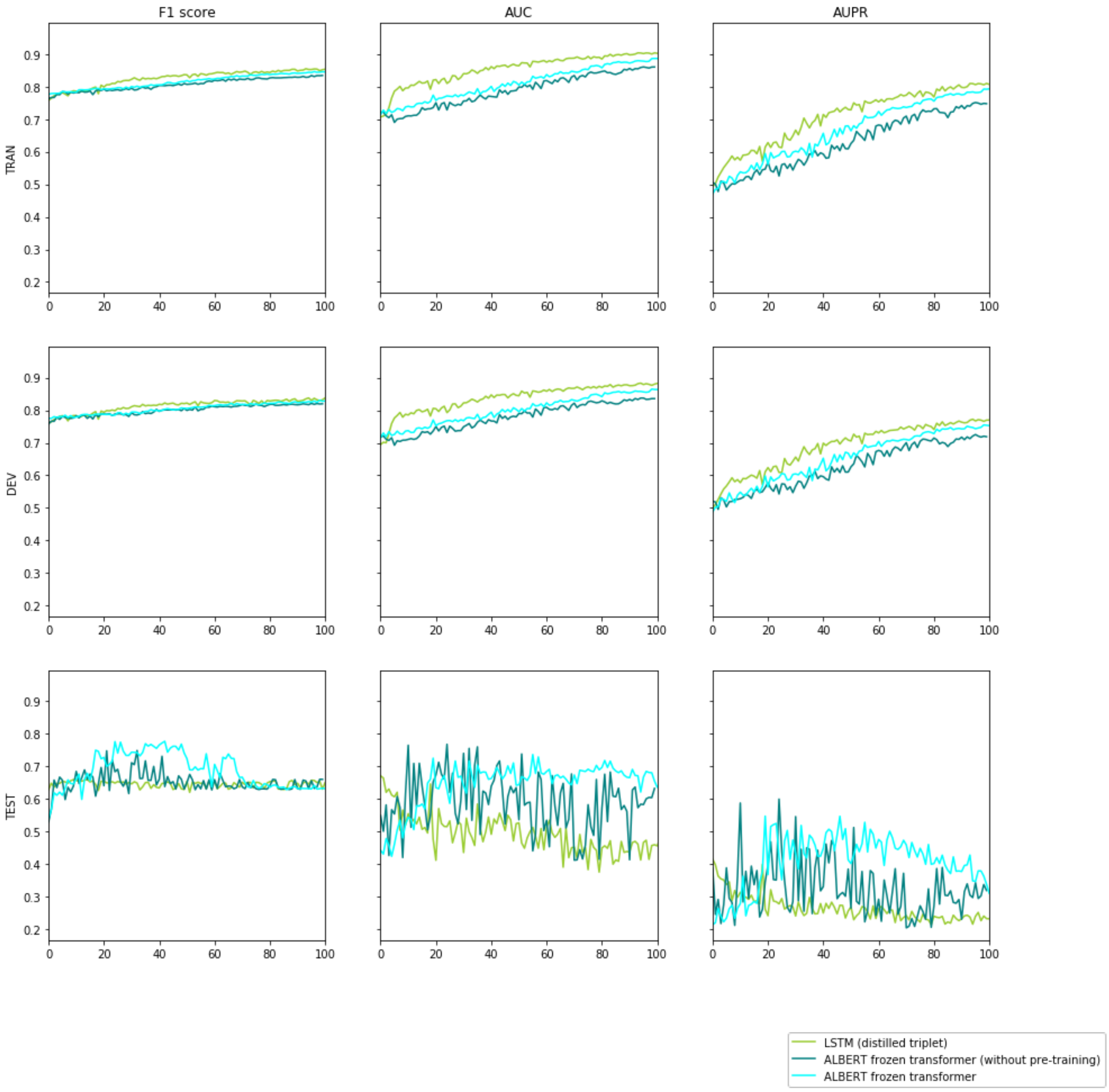


Supplementary Figure 4: The three major models as in the AUC curves in the paper. All x axis are the number of epochs trained. ALBERT frozen transformer proves consistent better performance, which becomes the critical advantage of DISAE in deorphanization, whhere remote orphan proteins are significantly different from training data and overfitting is impossible to control given the unknown true labels for classification. Using DISAE, we could be confident that even we train the model for too long or too short epochs, the prediction reliability on orphans will be robust.

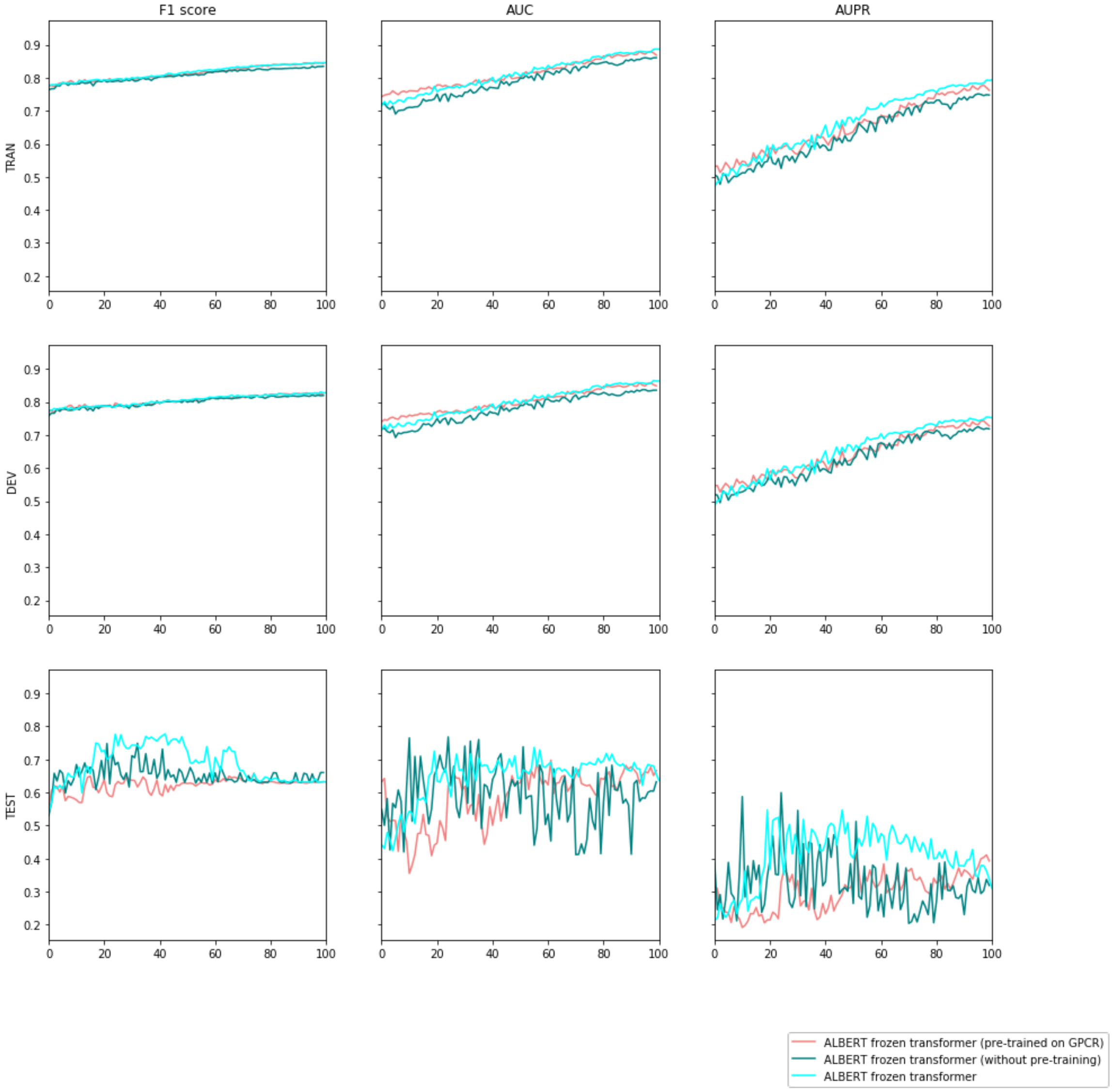


Supplementary Figure 5: The effect of pre-training: ALBERT frozen transformer, i.e. the one pre-trained on whole pfams as proposed in DISAE, show robust and consistent better performance. All x axis are the number of epochs trained.

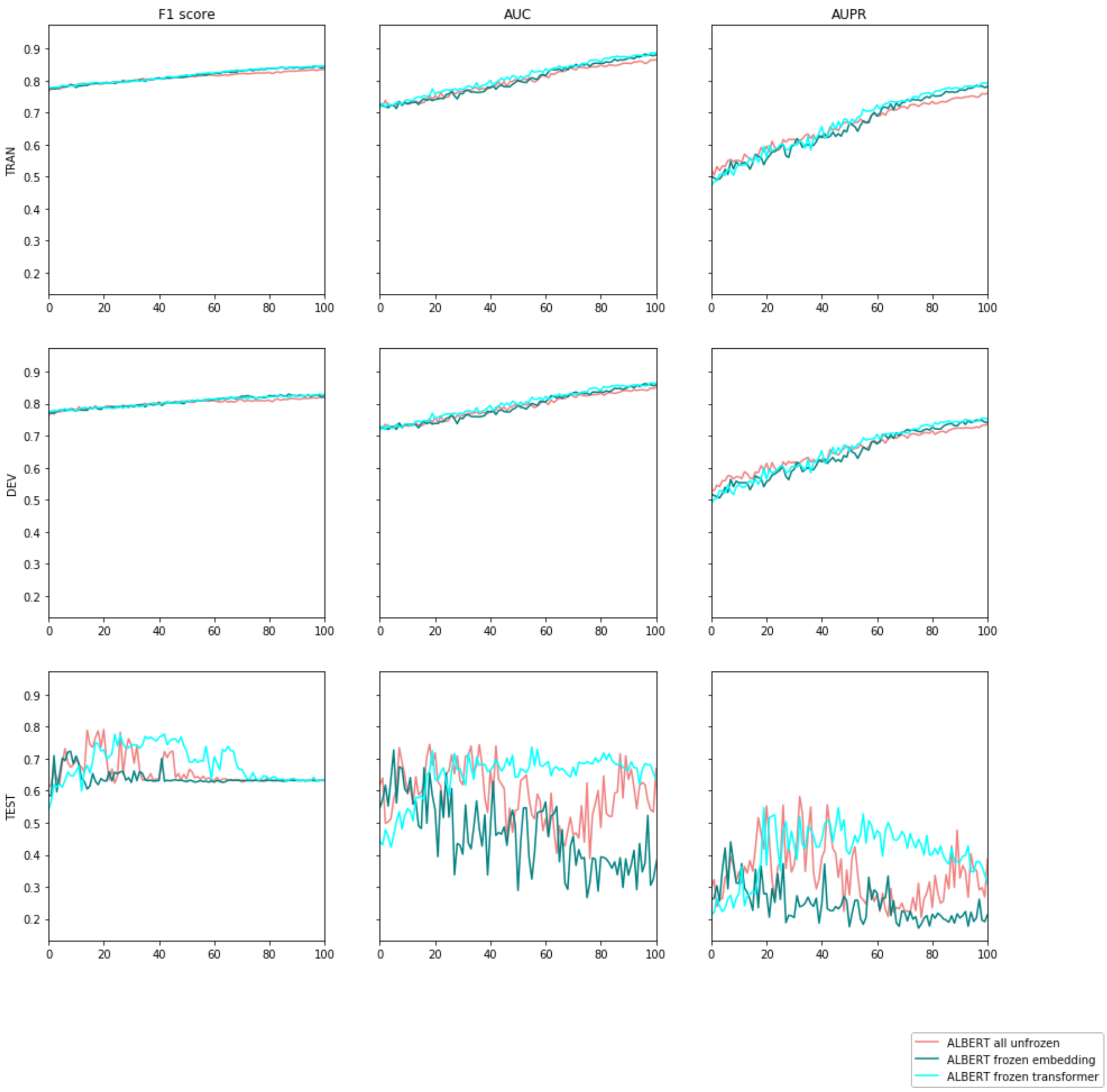


Supplementary Figure 6: The effect of different fine-tuning strategy with freezing parts of ALBERT: Although the other setting could be as good as ALBERT frozen transformer in some epochs, but they all suffer from large performance variance by epoch. ALBERT frozen transformer proves consistent and robust high performance. All x axis are the number of epochs trained

PF03402

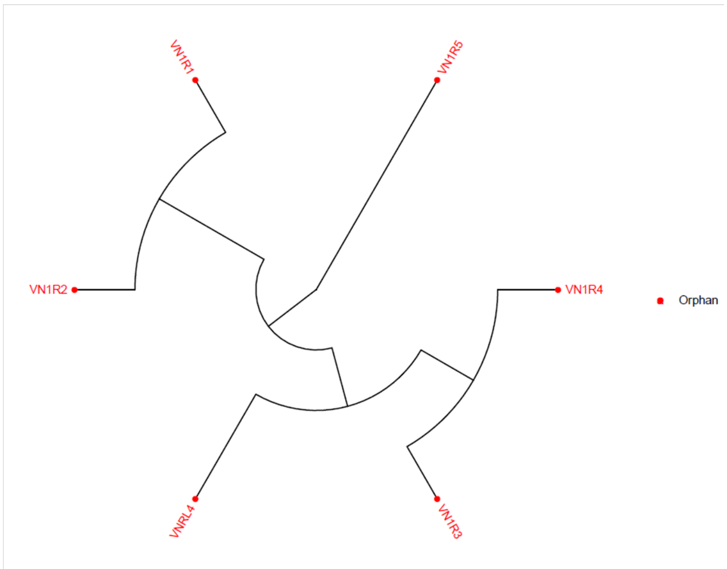


Supplementary Figure 7: Phylogenetic tree for PF03402

PF13853

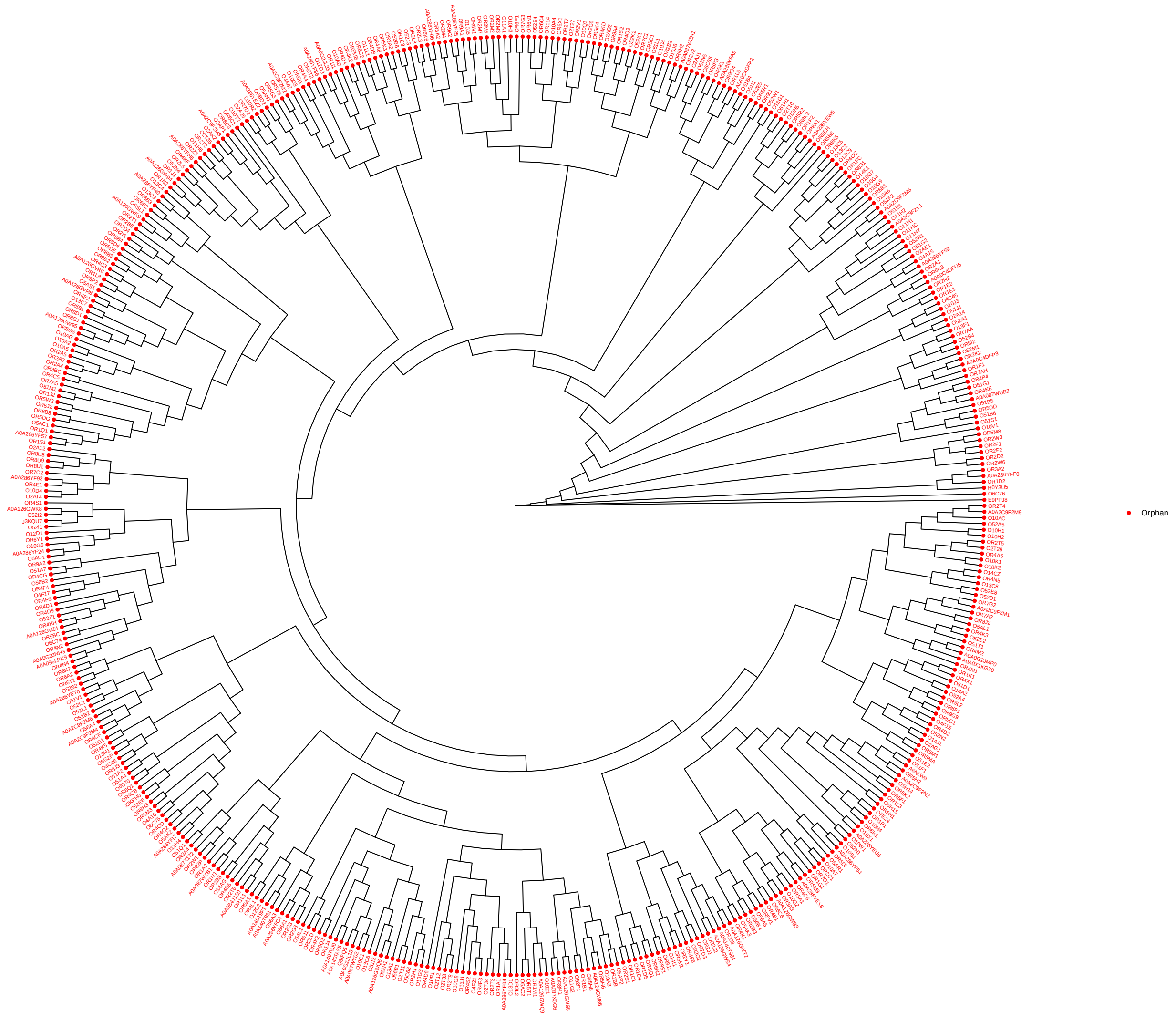


Supplementary Figure 8: Phylogenetic tree for PF13853

| | | ALBERT MODEL CONFIGURATION | |
|---|---|---|---|
| | | TRIPLETS FORM | SINGLET FORM |
| pre-training related | "attention_probs_dropout_prob" | 0 | |
| | "hidden_act": "gelu" | "gelu" | |
| | "hidden_dropout_prob" | 0 | |
| | "embedding_size" | 128 | |
| | "hidden_size" | 312 | |
| | "initializer_range" | 0.02 | |
| | "intermediate_size" | 1248 | |
| | "max_position_embeddings" | 512 | |
| | "num_attention_heads" | 12 | |
| | "num_hidden_layers" | 4 | |
| | "num_hidden_groups" | 1 | |
| | "net_structure_type" | 0 | |
| | "gap_size" | 0 | |
| | "num_memory_blocks" | 0 | |
| | "inner_group_num" | 1 | |
| | "down_scale_factor" | 1 | |
| | "type_vocab_size" | 2 | |
| | "vocab_size" | 19686 | 32 |
| | "ln_type" | / | "postln" |
| fine-tuning related | sequence embedding hidden units | 256 | |
| | protein sequence post-tokenization length | 210 | |

Supplementary Table 1: ALBERT configuration. ALBERT is using the package *Transformers* by Huggingface (https://github.com/huggingface/transformers). The author installed the package in Jan 2020.

| Neural-fingerprint CONFIGURATION | |
|---|---|
| Dropout | 0.1 |
| converlution layer size | 20 |
| converlution layer number | 4 |
| hidden units | 128 |
| Atomic connectivity degrees for chemical molecules | [0,1,2,3,4,5] |
| INERACTION PREDICTION CONFIGURATION | |
| design | two linear layer with batch normlization and ReLU |
| attentive pooling drop out | 0.3 |
| attentive pooling hidden units | 64 |
| LSTM CONFIGURATION | |
| embedding hidden units | 128 |
| number of LSTM layers | 1 |
| dropout | 0.2 |

Supplementary Table 2: There are three major components of the fine tuning model architecture: ALBERT to extract protein embedding, Neural-fingerprint to extract chemical embedding, and interaction prediction layers. LSTM serves as the baseline.

| OPTIMIZATION CONFIGURATION | |
|---|---|
| training epochs | 100 |
| batch size | 64 |
| optimizer | Adam |
| scheduler to adjust learning rate | cosineannealing |
| initial learnin rate | 2.00E-05 |
| L2 regularization weight | 1.00E-04 |
| Deep Learning Server | Supermicro SuperServer 4028GR-TR |
| GPU | NVIDIA Tesla® V100 with 32 GB per GPU (256 GB total) of GPU memory |
| CPU | Intel Xeon E5-2650 v4 2.2 GHz 12-Core (48-core total) |

Supplementary Table 3: Optimization related configuration

| Orphan Receptor | Predicted Drug [Known binding protein] |
|---|---|
| A0A087WY02 | XXPANQJNYNUNES-UHFFFAOYSA-N ['P23975', 'Q05940', 'P31645'] |
| A0A0A0MQW8 | BYJAVTDNIXVSPW-UHFFFAOYSA-N ['Q96RJ0', 'P35348', 'P43140'] |
| A0A0B4J1V8 | RGCVKNLCSQQDEP-UHFFFAOYSA-N ['P33261', 'P18901', 'P08684'] |
| A0A0C4DFX5 | MBUVEWMHONZEQD-UHFFFAOYSA-N ['P11509', 'P33261', 'Q16873'] |
| A0A126GVR8 | BPZSYCZIITTYBL-YJYMSZOUSA-N ['P13945', 'P11509', 'P07550'] |
| A0A126GWK9 | VQODGRNSFPNSQE-DVTGEIKXSA-N ['P46721', 'P04083', 'P20309'] |
| A0A126GWS4 | VMWNQDUVQKEIOC-CYBMUJFWSA-N ['P18901', 'P08684', 'P50226'] |
| A0A1B0GTK7 | ATALOFNDEOCMKK-OITMNORJSA-N ['P29371', 'P11712', 'P33261'] |
| A0A1B0GVZ0 | OWQUZNMMYNAXSL-UHFFFAOYSA-N ['Q01959', 'P31390', 'P35367'] |
| A0A286YF86 | BGDKAVGWHJFAGW-UHFFFAOYSA-N ['P08485', 'P20309', 'P17200'] |
| A0A286YF92 | IRSCQMHQWWYFCW-UHFFFAOYSA-N ['', 'Q13255', 'Q96FL8'] |
| A0A286YFH6 | BUGYDGFZZOZRHP-UHFFFAOYSA-N ['O00591', 'P11509', 'Q96FL8'] |
| A0A2C9F2M5 | CYQFCXCEBYINGO-IAGOWNOFSA-N ['P47746', 'P47936', 'P11712'] |
| A3KFT3 | KKZJGLLVHKMTCM-UHFFFAOYSA-N ['', 'Q16678', 'P33527'] |
| A6NFC9 | BYJAVTDNIXVSPW-UHFFFAOYSA-N ['Q96RJ0', 'P35348', 'P43140'] |
| A6NH00 | BGDKAVGWHJFAGW-UHFFFAOYSA-N ['P08485', 'P20309', 'P17200'] |
| A6NMS3 | IYIKLHRQXLHMJQ-UHFFFAOYSA-N ['P11509', 'P07550', 'P51589'] |
| A6NMU1 | BARDROPHSZEBKC-OITMNORJSA-N ['P08684', 'P11712', 'P25103'] |
| C9J1J7 | YFGHCGITMMYXAQ-LJQANCHMSA-N ['P20815', 'P11712', 'P05177'] |
| C9JQD8 | LWAFSWPYPHEXKX-UHFFFAOYSA-N ['P10635', 'P07550', 'P08588'] |
| C9JW47 | BYBLEWFAAKGYCD-UHFFFAOYSA-N ['Q6PIU1', 'P22001', 'P33261'] |
| E7ENI1 | OCJYIGYOJCODJL-UHFFFAOYSA-N ['P10635', 'P02768', 'P35367'] |
| E9PH76 | ZSCDBOWYZJWBIY-UHFFFAOYSA-N ['P07550', 'P33261', 'P11229'] |
| E9PPJ8 | OZYUPQUCAUTOBP-QXAKKESOSA-N ['P33535', 'P35372'] |
| F8VUV1 | CYQFCXCEBYINGO-IAGOWNOFSA-N ['P47746', 'P47936', 'P11712'] |
| G3V4Q5 | XXPANQJNYNUNES-UHFFFAOYSA-N ['P23975', 'Q05940', 'P31645'] |
| H0Y622 | UCHDWCPVSPXUMX-TZIWLTJVSA-N ['P11509', 'P07550', 'P21452'] |
| J3KQU7 | IZTQOLKUZKXIRV-YRVFCXMDSA-N ['Q9NPD5', 'P30553', 'Q63931'] |
| O00590 | JOATXPAWOHTVSZ-UHFFFAOYSA-N ['P13945', 'P07550', 'P18089'] |
| O15218 | BYJAVTDNIXVSPW-UHFFFAOYSA-N ['Q96RJ0', 'P35348', 'P43140'] |
| O43869 | UCHDWCPVSPXUMX-TZIWLTJVSA-N ['P11509', 'P07550', 'P21452'] |
| O60431 | XXPANQJNYNUNES-UHFFFAOYSA-N ['P23975', 'Q05940', 'P31645'] |
| O76099 | SHGAZHPCJJPHSC-YCNIQYBTSA-N ['P31025', 'P11509', 'Q02928'] |
| P04000 | LUZRJRNZXALNLM-JGRZULCMSA-N ['P08684', 'P34969', 'P35462'] |
| P04001 | XXPANQJNYNUNES-UHFFFAOYSA-N ['P23975', 'Q05940', 'P31645'] |
| P0C604 | DGBIGWXXNGSACT-UHFFFAOYSA-N ['O00591', 'O14764', 'P48169'] |
| P0C628 | HTIQEAQVCYTUBX-UHFFFAOYSA-N ['Q02641', 'P08684', 'P04798'] |
| P0C645 | IRSCQMHQWWYFCW-UHFFFAOYSA-N ['', 'Q13255', 'Q96FL8'] |
| P0C7N5 | VMWNQDUVQKEIOC-CYBMUJFWSA-N ['P18901', 'P08684', 'P50226'] |
| P0DMS8 | DGBIGWXXNGSACT-UHFFFAOYSA-N ['O00591', 'O14764', 'P48169'] |
| P0DN77 | XXPANQJNYNUNES-UHFFFAOYSA-N ['P23975', 'Q05940', 'P31645'] |
| P0DN78 | XXPANQJNYNUNES-UHFFFAOYSA-N ['P23975', 'Q05940', 'P31645'] |
| P47804 | QWAXKHKRTORLEM-UGJKXSETSA-N ['P21452', 'P08684', 'P35462'] |
| P58173 | VHYCDWMUTMEGQY-UHFFFAOYSA-N ['P13945', 'P07550', 'P08588'] |
| Q15612 | OIRDTQYFTABQOQ-KQYNXXCUSA-N ['P29275', 'P55263', 'P25099'] |
| Q15617 | ZSCDBOWYZJWBIY-UHFFFAOYSA-N ['P07550', 'P33261', 'P11229'] |
| Q6IFH4 | KWTSXDURSIMDCE-QMMMGPOBSA-N ['P25100', 'Q8HZ64', 'P10635'] |
| Q7Z5H5 | IYIKLHRQXLHMJQ-UHFFFAOYSA-N ['P11509', 'P07550', 'P51589'] |
| Q8N0Y5 | COUYJEVMBVSIHV-SFHVURJKSA-N ['P10632', 'P07550', 'P11712'] |
| Q8N6U8 | BYJAVTDNIXVSPW-UHFFFAOYSA-N ['Q96RJ0', 'P35348', 'P43140'] |
| Q8NG76 | IZTQOLKUZKXIRV-YRVFCXMDSA-N ['Q9NPD5', 'P30553', 'Q63931'] |
| Q8NG83 | CJOFXWAVKWHTFT-XSFVSMFZSA-N ['Q9HB55', 'P22086', 'P10635'] |
| Q8NG85 | IYIKLHRQXLHMJQ-UHFFFAOYSA-N ['P11509', 'P07550', 'P51589'] |
| Q8NGB2 | BARDROPHSZEBKC-OITMNORJSA-N ['P08684', 'P11712', 'P25103'] |
| Q8NGC2 | MEFKEPWMEQBLKI-AIRLBKTGSA-N ['P19623', 'P31153', 'P25099'] |
| Q8NGC8 | COUYJEVMBVSIHV-SFHVURJKSA-N ['P10632', 'P07550', 'P11712'] |
| Q8NGD3 | OCJYIGYOJCODJL-UHFFFAOYSA-N ['P10635', 'P02768', 'P35367'] |

**Table 4 continued from previous page**

| Orphan Receptor | Predicted Drug [Known binding protein] |
|---|---|
| Q8NGD4 | UCTWMZQNUQWSLP-VIFPVBQESA-N ['P07550', 'O15244', 'P08684'] |
| Q8NGE9 | WNTYBHLDCKXEOT-UHFFFAOYSA-N ['P14416', 'P31388', 'P10275'] |
| Q8NGG2 | BGDKAVGWHJFAGW-UHFFFAOYSA-N ['P08485', 'P20309', 'P17200'] |
| Q8NGI3 | CYQFCXCEBYINGO-IAGOWNOFSA-N ['P47746', 'P47936', 'P11712'] |
| Q8NGI4 | IRSCQMHQWWYFCW-UHFFFAOYSA-N ['', 'Q13255', 'Q96FL8'] |
| Q8NGI7 | OCJYIGYOJCODJL-UHFFFAOYSA-N ['P10635', 'P02768', 'P35367'] |
| Q8NGJ5 | JSWZEAMFRNKZNL-UHFFFAOYSA-N ['P11712', 'P41595', 'P05177'] |
| Q8NGJ9 | GEFQWZLICWMTKF-CDUCUWFYSA-N ['P25100', 'P18089', 'P08913'] |
| Q8NGK2 | LUZRJRNZXALNLM-JGRZULCMSA-N ['P08684', 'P34969', 'P35462'] |
| Q8NGK6 | ZSCDBOWYZJWBIY-UHFFFAOYSA-N ['P07550', 'P33261', 'P11229'] |
| Q8NGL9 | UCHDWCPVSPXUMX-TZIWLTJVSA-N ['P11509', 'P07550', 'P21452'] |
| Q8NGM9 | BYJAVTDNIXVSPW-UHFFFAOYSA-N ['Q96RJ0', 'P35348', 'P43140'] |
| Q8NGN7 | XXPANQJNYNUNES-UHFFFAOYSA-N ['P23975', 'Q05940', 'P31645'] |
| Q8NGR4 | BGDKAVGWHJFAGW-UHFFFAOYSA-N ['P08485', 'P20309', 'P17200'] |
| Q8NGS1 | XXPANQJNYNUNES-UHFFFAOYSA-N ['P23975', 'Q05940', 'P31645'] |
| Q8NGT5 | ATALOFNDEOCMKK-OITMNORJSA-N ['P29371', 'P11712', 'P33261'] |
| Q8NGU2 | UCHDWCPVSPXUMX-TZIWLTJVSA-N ['P11509', 'P07550', 'P21452'] |
| Q8NGW1 | DRHKJLXJIQTDTD-OAHLLOKOSA-N ['P08684', 'P34969', 'P35462'] |
| Q8NGX9 | URKOMYMAXPYINW-UHFFFAOYSA-N ['P31389', 'P07550', 'P33261'] |
| Q8NGY9 | UCTWMZQNUQWSLP-VIFPVBQESA-N ['P07550', 'O15244', 'P08684'] |
| Q8NGZ2 | IZTQOLKUZKXIRV-YRVFCXMDSA-N ['Q9NPD5', 'P30553', 'Q63931'] |
| Q8NGZ4 | RUDATBOHQWOJDD-BSWAIDMHSA-N ['P52895', 'Q96RI1', 'P08684'] |
| Q8NH04 | BYBLEWFAAKGYCD-UHFFFAOYSA-N ['Q6PIU1', 'P22001', 'P33261'] |
| Q8NH05 | KKGQTZUTZRNORY-UHFFFAOYSA-N ['P21453', 'O95977', 'P43004'] |
| Q8NH41 | OZVBMTJYIDMWIL-AYFBDAFISA-N ['P07550', 'P08684', 'P34969'] |
| Q8NH48 | ZSCDBOWYZJWBIY-UHFFFAOYSA-N ['P07550', 'P33261', 'P11229'] |
| Q8NH51 | BUGYDGFZZOZRHP-UHFFFAOYSA-N ['O00591', 'P11509', 'Q96FL8'] |
| Q8NH57 | XXPANQJNYNUNES-UHFFFAOYSA-N ['P23975', 'Q05940', 'P31645'] |
| Q8NH61 | RUDATBOHQWOJDD-BSWAIDMHSA-N ['P52895', 'Q96RI1', 'P08684'] |
| Q8NH74 | BYJAVTDNIXVSPW-UHFFFAOYSA-N ['Q96RJ0', 'P35348', 'P43140'] |
| Q8NH87 | GJPICJJJRGTNOD-UHFFFAOYSA-N ['P25101', 'P11712', 'O95342'] |
| Q8NH89 | ZSCDBOWYZJWBIY-UHFFFAOYSA-N ['P07550', 'P33261', 'P11229'] |
| Q8NH95 | DERZBLKQOCDDDZ-JLHYYAGUSA-N ['P11509', 'O60840', 'P11229'] |
| Q8TCB6 | HTIQEAQVCYTUBX-UHFFFAOYSA-N ['Q02641', 'P08684', 'P04798'] |
| Q8TDV2 | XXPANQJNYNUNES-UHFFFAOYSA-N ['P23975', 'Q05940', 'P31645'] |
| Q8WZA6 | ZFXYFBGIUFBOJW-UHFFFAOYSA-N ['P29275', 'P25099', 'Q60614'] |
| Q96KK4 | XXPANQJNYNUNES-UHFFFAOYSA-N ['P23975', 'Q05940', 'P31645'] |
| Q96R54 | VMWNQDUVQKEIOC-CYBMUJFWSA-N ['P18901', 'P08684', 'P50226'] |
| Q96R67 | ZSCDBOWYZJWBIY-UHFFFAOYSA-N ['P07550', 'P33261', 'P11229'] |
| Q9BZJ7 | UCHDWCPVSPXUMX-TZIWLTJVSA-N ['P11509', 'P07550', 'P21452'] |
| Q9BZJ8 | KWTSXDURSIMDCE-QMMMGPOBSA-N ['P25100', 'Q8HZ64', 'P10635'] |
| Q9GZM6 | PVNIIMVLHYAWGP-UHFFFAOYSA-N ['P49019', 'Q15274', 'Q80Z39'] |
| Q9H1Y3 | FIVSJYGQAIEMOC-ZGNKEGEESA-N ['P10635', 'Q9UNQ0', 'P25103'] |
| Q9H210 | IYIKLHRQXLHMJQ-UHFFFAOYSA-N ['P11509', 'P07550', 'P51589'] |
| Q9H339 | IYIKLHRQXLHMJQ-UHFFFAOYSA-N ['P11509', 'P07550', 'P51589'] |
| Q9NZP2 | IQVRBWUUXZMOPW-PKNBQFBNSA-N ['P29275', 'Q96FL8', 'P25099'] |
| Q9UGF5 | GHOSNRCGJFBJIB-UHFFFAOYSA-N ['P30556', 'P11712', 'P23219'] |
| Q9UGF6 | BTCSSZJGUNDROE-UHFFFAOYSA-N ['Q7Z2H8', 'Q9Z0U4', 'Q9UBS5'] |
| Q9UHM6 | BGDKAVGWHJFAGW-UHFFFAOYSA-N ['P08485', 'P20309', 'P17200'] |

Supplementary Table 4: Predicted approved drug examples: 649 human orphan GPCRs, each paired to 555 approved GPCR-targeted drugs as novel samples chemical-protein pairs. Listed here are 106 of the orphan proteins paired with at least one approved GPCR-targeted drugs with estimated false positive rate lower than 0.05. Proteins are presented with Uniprot Id and chemicals are presented with InChIKey.
